# Site-suitability difference and the expectation of species richness difference in stochastic two-subcommunities models

**DOI:** 10.1101/2023.01.01.522407

**Authors:** Ryosuke Iritani

## Abstract

Understanding how community assembly during the initial faunization phases determines the difference in species richness is a fundamental question in ecology. However, there are few predictions for when and to what degree the differences in species richness between subcommunities will emerge. Here, we investigate the expectation of richness difference in a pair of subcommunities, assuming that species may have different and independent presence probabilities. We then introduce several indices and examine how expected richness difference is determined by the indices. We found that (i) species differences (the average of species presence probabilities in two subcommunities divided by the total presence probability) have inconsistent effects on richness difference; (ii) the degree of spatial heterogeneity (average of differences in species presence probabilities in two subcommunities across species) may, but not always, have a good predictive ability for richness difference; and (iii) the absolute difference in average presence probabilities (site-suitability difference) has robust predictive ability for richness difference unless richness difference is very small. This work provides a theoretical framework for predicting and analyzing species richness difference from presence-absence data based on null models.

## 1 Introduction

Why, and how much, does one area have richer species number than does another? Addressing this question and predicting the patterns of richness difference are ultimate goals in ecology [1–3]. Richness difference (the absolute difference between two local species-richnesses) is caused by various assemblage processes such as distance, area, species differences in colonization ability, and environmental filtering [4–7]. Knowledge in richness differences provides informative facets, for example, for conservation planning of biodiversity hotspots [8, 9], the global pattern of richness gradient [10–12], and spatial variations in biodiversity (beta-diversity; [13, 14]). It is thus of great importance to predict how richness difference is determined by various biotic and abiotic factors.

A challenging problem in predicting richness difference results from heterogeneous factors in natural communities [15–18]. Such factors include distance, degree of fragmentation, disturbance, and colonization ability [11, 12]. Surprisingly few theories exist to make predictions for richness difference. Iritani *et al*. [19], by handling species-incidence (presence or absence) as a probabilistic variable [2], studied how species difference and spatial heterogeneity affect beta-diversity, but did not consider richness difference. Similarly, based on the assumption that species incidence is independent, Calabrese *et al*. [20] and Carmona & Pärtel [21] took the probabilistic approach to estimate the species richness and its stochastic variations. However, neither did their studies consider richness difference. Taken together, the current methodological tools have not yet succeeded in providing theoretical predictions of how richness difference can be shaped by stochastic processes when there are species difference and spatial heterogeneity.

In this article, we derive an exact, analytical formula for richness difference between two subcommunities, assuming that: (i) species incidence is independent across space and species, and (ii) presence probabilities of species in two subcommunities are allowed to take differential values. Then we use three predictive indices (predictors) and examine their predictive abilities for richness difference. The present theoretical work provides useful guide to predict how richness difference can occur in natural and experimental environments, especially by nullifying correlations among species incidence.

## 2 Method and result

### 2.1 General analysis

We write *x*_*i,j*_ for the binary state of species *i*’s incidence in site *j* (*p*_*i,j*_ = 1 if *i* is present in *j*, otherwise absent *p*_*i,j*_ = 0), and *p*_*i,j*_ for the probability that species *i* is present in site *j*. The probabilities *p*_*i,j*_ may be, for example, estimated from long time-series data of species incidence (e.g., BioTIME [22]). We assume that a species-pool has a limited size, such that at most *S* species can be present in at least one of the sites (more technically, the species pool consists of species *i* that can inhabit at least one of the sites with a strictly positive probability, min {*p*_*i*,1_, *p*_*i*,2_} > 0). The symbols are summarized in Appendix A.

The species richness, or alpha-diversity, in site *j* is given by:

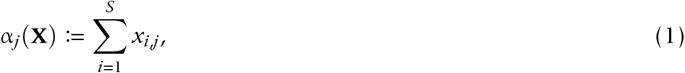

which is a nonnegative integer value, *α_j_* = 0, 1, …, *S*. In statistics, *α_j_* is said to follow Poisson-Binomial distribution with parameters **p**_*j*_ ≔ (*p*_1,*j*_, *p*_2,*j*_, …, *p*_*S,j*_) [23]. The expected value of alpha-diversity is given by:

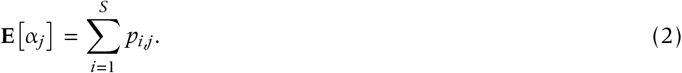

Denoting by Λ(**X**) ≔ min {*α*_1_ (**X**), *α*_2_ (**X**)} the minimum of the species richness, we can write down its expected value, **E**[Λ], in an analytically closed form (Appendix B). Using **E**[Λ] and writing *D*(**X**) ≔ |*α*_1_(**X**) − *α*_2_(**X**)| for the difference of species richness between two subcommunities, we use the identity |*α*_1_ − *α*_2_| ≡ *α*_1_ + *α*_2_ − 2 min {*α*_1_, *α*_2_} to get:

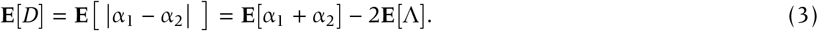

### 2.2 Predictors

We use three parameters to characterize the variations in the presence probabilities, *p*_*i,j*_. First is species difference denoted *w*, which is defined as the proportion of the average taken for two sites of absolute deviation of the presence probabilities from within-site average presence probability to the sum of presence probabilities for all species for two sites [19]. Second is spatial heterogeneity denoted *h*, which is defined as the average, taken across species, of the difference of presence probabilities for two sites. These are given by:

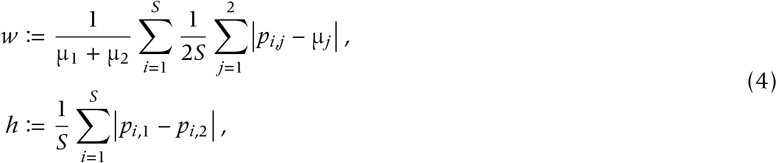

where 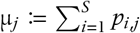 represents the average presence probability of species in site *j* [19]. Species difference is zero *w* = 0 if and only if all species are identical in the presence probabilities. Finally, we propose the following quantity (“site-suitability difference”):

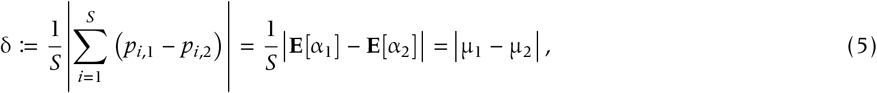

which represents the absolute difference in average presence probabilities in sites 1 and 2. Note that δ ≤ *h* by the triangle inequality (in which the equality, *h* = δ, occurs when one site is predominantly more suitable than the other: *p*_*i*,1_ ≥ *p*_*i*,2_ for all *i* = 1, …, *S* or either *p*_*i*,1_ ≤ *p*_*i*,2_ for all *i* = 1, …, *S*), and also that *S* ∙ δ ≤ **E**[*D*] by Jensen’s inequality [24].

### 2.3 Simulations

To examine how richness difference depends on these three indices, we carried out stochastic simulations based on the following scheme: (1) setting *S* = 16, we choose a pair of probability distributions to generate **p**_*j*_ = *p*_1,*j*_, …, *p*_*S,j*_ for site 1 and 2; we refer to such distributions as ‘incidence-generators’ in this article. The number of simulations is 300. (2) We compute the species difference, spatial heterogeneity, site-suitability difference, and expected richness difference, and then plot scatter plots.

We illustrate the theoretical result using three sets of the beta distributions for incidence-generators (and we write “LHS ~ Beta[•,•]” to mean that LHS follows the Beta distribution with parameters indicated by [•, •]).

1. “Random model”: *p*_*i*,1_ ~ Beta[1.10, 1.09] and *p*_*i*,2_ ~ Beta[1.09, 1.10];
2. “Overlap model”: *p*_*i*,1_ ~ Beta[1.28,0.96] and *p*_*i*,2_ ~ Beta[0.96,1.28]; and
3. “Dominance model”: *p*_*i*,1_ ~ Beta[1.6,0.4] and *p*_*i*,2_ ~ Beta[0.4,1.6].

For the random model, in which there is no clear small-large relationship between *p*_*i*,1_ and *p*_*i*,2_, neither species difference nor spatial heterogeneity predict the richness difference accurately, but site-suitability difference has a monotonic, and very consistent, effect of richness difference (Figure 1).

**Figure 1:**
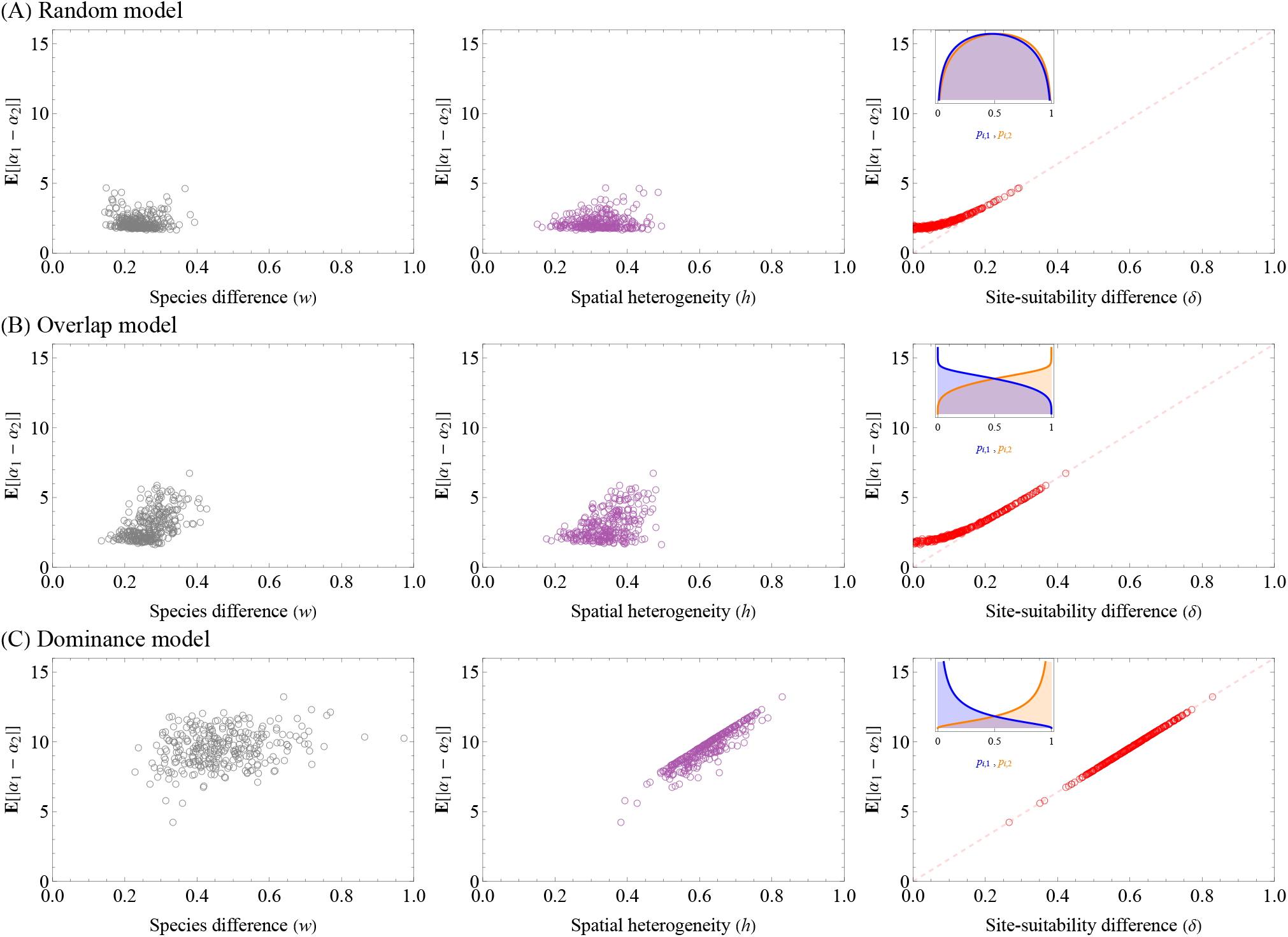
The performance of species difference (left), spatial heterogeneity (middle), and site-suitability difference (right), for predicting the expectation of richness difference, in three models producing different probability distributions of **p**_1_ and **p**_2_. In the right panels, the dotted segment is a line with slope *S*. Inset panels are the incidence-generators (i.e., the probability density functions) of **p**_1_ (blue) and **p**_2_ (orange). (A) When **p**_1_ and **p**_2_ are drawn from nearly indentical distributions (incidence-generator), richness difference is systematically small because the alpha diversity is likely similar between two sites. Neither can three predictors perform well, although site-suitability difference seems to have a formal relationship with richness difference. (B) When the two incidence-generators are partially (but not fully) overlapped, the predictive ability of site-suitability difference for species richness becomes better, whereas the other two indices do not show consistent patterns. (C) When one site is dominantly more likely to be inhabited by species than is the other, the predictive ability of site-suitability difference for the species becomes almost perfect. Parameters: *S* = 16; the number of simulations (i.e., circles) in each of panels (A–C): 300.

For the overlap model, in which *p*_*i*,2_ is likely to be larger than *p*_*i*,1_ compared to the random model case, richness difference has no clear dependence on species difference. The predictive abilities of both spatial heterogeneity and site-suitability for richness difference are improved over the random model case (Figure 1A and B), but the site-suitability’s predictive ability decreases as site-suitability and richness difference are smaller (Figure 1B).

For the dominance model, in which *p*_*i*,2_ is very likely to be larger than *p*_*i*,1_, we find that whereas species difference have a low predictive ability, spatial heterogeneity and site-suitability difference both almost perfectly (and linearly) predict richness difference; it should be however noted that this coincidence is in part due to the equality δ = *h* for the dominance model (Figure 1C). We found that these predictions are robust against changing the incidence-generators (the uniform, instead of beta distributions) as well as limited observations of incidence-tables (Appendix C).

In sum, we examined how accurately species difference index, spatial heterogeneity index, and site-suitability difference index can predict the expectation of richness difference. The predictive abilities of these indices strongly depend on the nature of probability distributions for species incidence. When species’ presence probabilities in two sites are very similar as in the random model, it is difficult to predict the richness difference, but there seems to be a governing relation of site-suitability difference with richness difference. As the probability distributions for the two sites become dissimilar (from random, overlap, to dominance model), the predictive abilities of the site-suitability and spatial heterogeneity are improved.

## 3 Discussion

We investigated how accurately the expectation of richness difference can be predicted by three indices. These indices can be estimated from long time-series data of species incidence. We found that the expectation of richness difference has a complicated, and almost intractable, analytical form. While species differences and site-heterogeneity both have limited predictive ability of the expected richness difference, a site-suitability index, defined as the average of the absolute difference of alpha diversity between two sites, provides a good predictor. However, the predictive ability of the index decreases as the site-suitability index becomes small. The dependence of richness difference on site-suitability difference seems to be positive and nonlinear (though asymptotically linear).

The predictive ability of the spatial heterogeneity for richness difference is sensitive to the probability distributions with which species are present in each site. Specifically, if one site is a priori known to have predominantly larger species richness than does the other site, then spatial heterogeneity estimated from long-time series data of species presence-absence can be used as a measure of richness difference. On the other hand, without such a priori knowledge, i.e., if the average probabilities of species to be present in one site and in the other are similar, it would be difficult to statistically infer richness difference from site-heterogeneity measure alone. This is because site-heterogeneity measure is invariant with the permutation of sites for each species: exchanging *p*_*i*,1_ and *p*_*i*,2_ for each species *i* does not change the value of *h*, and thus does not detect absolute richness difference.

In addition, our key finding predicts that richness difference may be adequately predicted by the site-suitability index, all things being equal. Biologically, despite the presence of heterogeneity, the absolute difference in alpha-diversity provides a good estimate for the expected absolute difference in species richness. However, the fit becomes less accurate when the site-suitability and richness difference are both low, suggesting that it is difficult to estimate the richness difference from empirical data that typically contains a limited number of unique species. We reason this result as follows. First of all, the absolute value and its original value are equivalent (namely|*x*| = *x*) whenever the latter *x* is positive (‘positivity invariance rule’). When site-suitability difference is large, it is immediately clear that one site (say site 2) tends to have predominantly higher alpha-diversity as does the other (site 1; dominance model case, Figure 1C). In this case, *α*_2_ − *α*_1_ is likely positive with a very high probability, so that the positivity-invariance rule applies and the original value provides the good estimate, although with some error associated with the case *α*_2_ < *α*_1_. In contrast, if site-suitability difference is small, then two sites are likely to have similar species richness (as in random model case; Figure 1A); in this case, the positivity-invariance rule does not apply because the sign of *α*_1_ − *α*_2_ varies. Biologically, site-suitability difference being zero likely occurs when two sites are homogeneous. Thus, heterogeneity may lower the predictive ability of site-suitability difference for richness difference.

Our model can be used as a framework of environmentally constrained null-model analyses of richness difference [15]. Despite the intuition that incorporating species differences and spatial heterogeneity matters, the present work showed that the effects of such heterogeneity are overall small unless the absolute difference of the expectations of alpha-diversity in two sites is low. Specifically, by averaging long time-series data of incidence (e.g., [22]; also see Appendix C3), one may estimate presence probabilities, assuming that presence probabilities are at dynamical equilibrium [2]. In this vein, deviations of observed richness difference from the expectation of the present work provide a signal of the effect of certain nullified factors. Since we assumed that species incidence are uncorrelated and also that two sites are uncorrelated, such deviation can be explained by these factors.

Our work has further implications for predicting beta-diversity, because richness difference is a subcomponent beta-diversity (spatial variations in species richness; [6, 25]). Specifically, Baselga [6] proposed to partition incidence-based beta-diversity measures, e.g., Jaccard or Sørensen’s [26–28], into additive components of richness difference and turnover. Iritani *et al*. [19] derived the exact formula for the expectation of Jaccard dissimilarity in the same setting as the current work’s, but further methodology has yet to be developed for the additive partitioning scheme proposed by Baselga [6, 25]. Future studies may apply the methodology to the additive partitioning frameworks and thereby advance the theory of beta-diversity for the heterogeneous world.

The present study has several avenues for generalizations, e.g., considering multiple-sites case, relaxing species-independence assumption, and analyzing abundance-based nestedness components. Future studies may aim to make testable predictions for, and improve our understanding of, the repeated patterns of spatial biodiversity [29].

In conclusion, we derived the analytical formula for the expected richness-difference based on the usual assumption that species incidences are uncorrelated across space. Our stochastic simulations showed that richness difference is very well explained by site-suitability, but the explanatory power is reduced when site-suitability index is small. Richness difference is a fundamental quantity and has the potential for future studies to produce interesting hypotheses of beta-diversity patterns.

## Acknowledgement and declaration

We thank Naoto Shinohara for helpful comments; and JSPS KAKENHI (grant numbers 19K22457, 19K23768 and 20K15882) for funding. We have no competing interests.

## Appendices

## A Notation

· *i*: label for species
· *S*: the maximum number of species that can be present in at least one of the sites
· *j*: label for site
· *x*_*i,j*_ : incidence of species *i* in site *j*
· **P**[proposition]: the probability that the proposition holds true
· **X**: matrix (table) with elements *x*_*i,j*_
· *p*_*i,j*_ : the probability of species *i* present in site *j*
· **E**[]: expectation over all possible **X**s
· 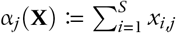: the species richness in site *j*
· 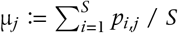: the average presence probability of speicies in site *j*
· *D*(**X**) ≔ |*α*_1_(**X**) − *α*_2_(**X**)|: the absolute difference of richness

## B Expectation of richness difference

In general, for any nonnegative integer-valued stochastic variable *z*, the following identity holds true:

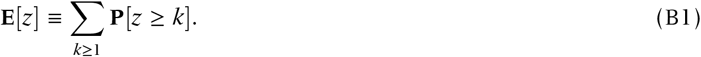

If we take *z* = min {*z*_1_, *z*_2_}, that is, if we aim to calculate the minimum of two independent random variables, then:

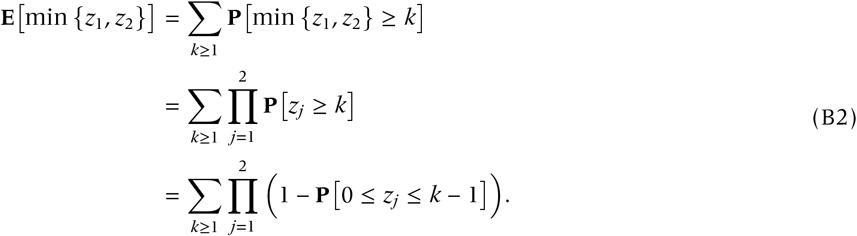

By taking *z_j_* = *α_j_*, the formula leads to:

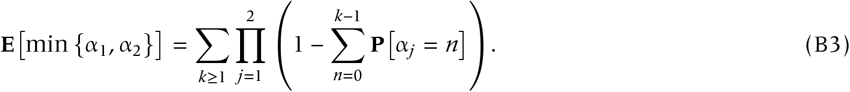

Since *α_j_* follows the Poisson-Binomial distribution (PBD) with parameters **p**_*j*_ (≔ (*p*_1,*j*_, …, *p*_*S,j*_),

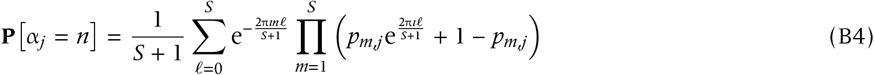

[23], where *ı* represents the imaginary unit. Therefore,

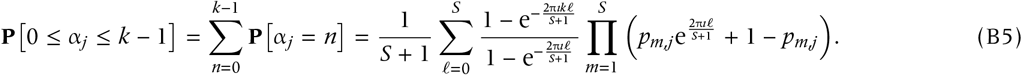

Note that the factor with *ℓ* = 0 in the summand is a removal singularity for the fraction. Inserting the last equation into Equation (B3) supplies the expectation of the minimum richness. Using the identity *α*_1_ +*α*_2_ − 2 min {*α*_1_, *α*_2_} ≡ |*α*_1_ − *α*_2_| and taking the expectation in both sides, we get the expectation of richness difference.

## C Robustness

## C.1 The uniform distributions

We choose the uniform distributions as the incidence-generators of **p_j_**. We found that the predictions obtained with the beta distributions are robust against this change (Figure SI-1).

## C.2 Averaging: Binomial approximation

As before we first generate **p**_*j*_ using the same scheme and then calculate μ_*j*_, and list it in a vector with length *S*: 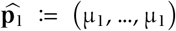 and 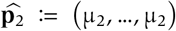. Then calculate the exact expectation **E**[*D*] (using **p**_1_ and **p**_2_) and the approximated expectation based on the approximation (using 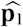 and 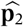). We found that the approximation tends to underestimate the true expectation, but the error becomes lower as the incidence-generators become more dissimilar (Figure SI-2).

## C.3 Averaging the raw values of δ and *D*

So far, we used the expectation of richness difference and site-suitability from *p*_*i,j*_*s*, and compared their values. In real situations, however, we estimate the mean richness difference and mean site-suitability both from incidence tables and compare them. That is, the comparison of the true expectations (**E**[*D*] versus δ) and that of means (average of *D* versus average of δ) are different.

Here we assess whether the difference is large. As before, we generate **p**_*j*_ from incidence-generators:

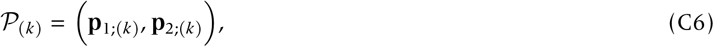

each of which is drawn from a fixed probability distribution (beta distribution), where *k* represents the simulation number, and incidence-tables (‘empirical tables’) from **p**_*j*_, with the number of the empirical tables, *N*, set to 300 (note that with *S* = 16 we have 2^32^ ≈ 10^10^ possible incidence-tables, far larger than 300):

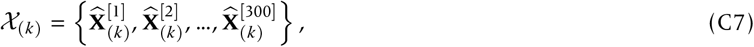

where each 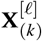 for *ℓ* = 1, …, 300 is drawn from the common, joint Beta distribution. We then compute (i) the true expectation of richness difference as well as (ii) the average of the absolute values of the richness difference for the empirical tables:

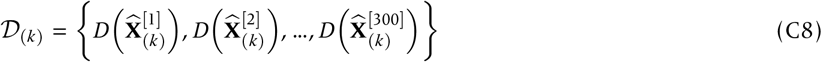

We then get:

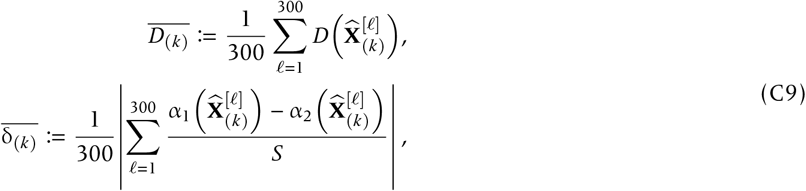

We repeat this procedure 200 times and compared 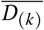 and 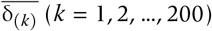 by depicting a scatter plot (Figure SI-3).

---

**SI Figure SI-1:**
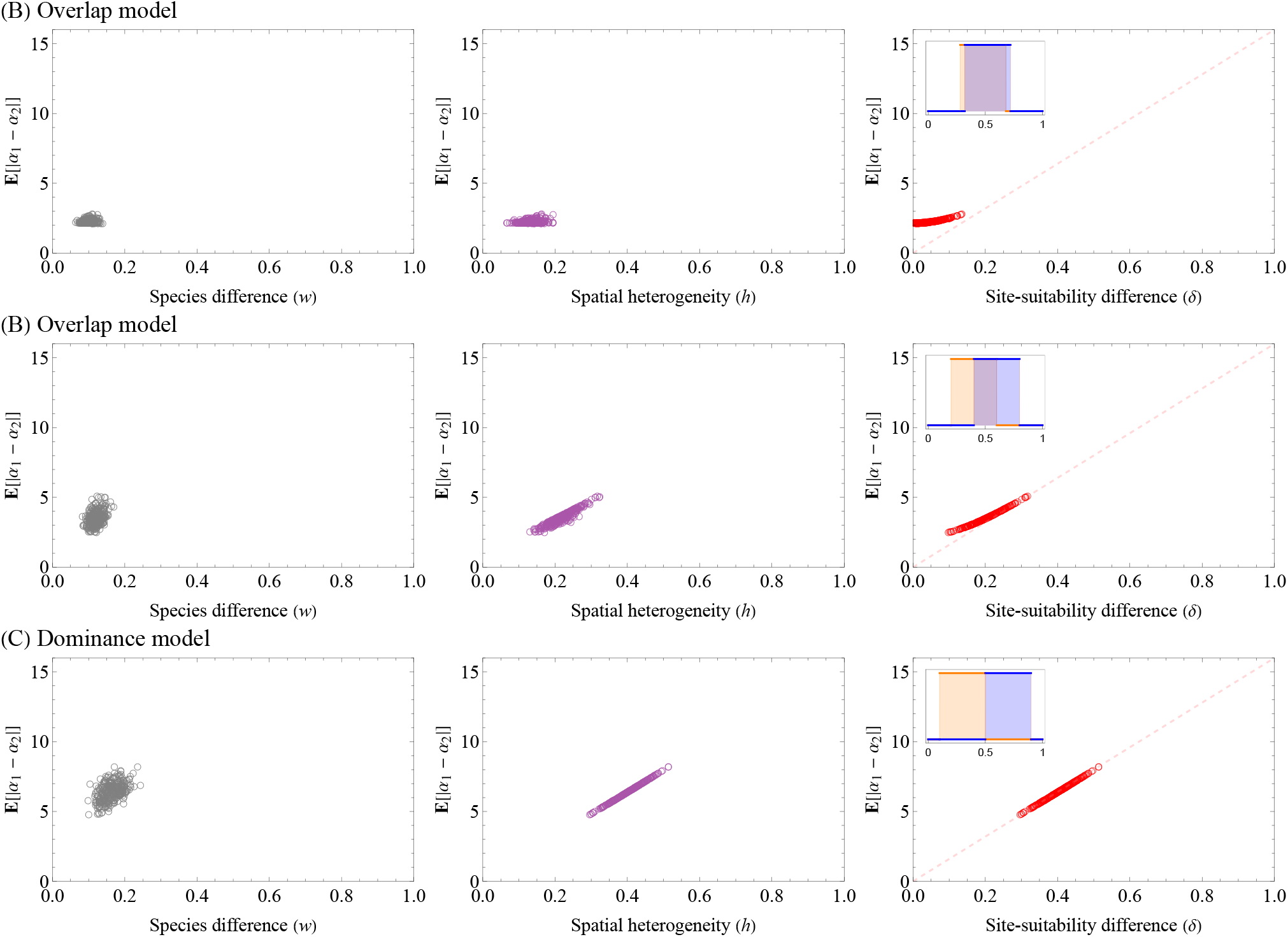
The same figure as the one in the main text, except the choice of the incidence-generators. Uniform distributions on: (A) [0.28, 0.68] and [0.32, 0.72]; (B) [0.2, 0.6] and [0.4, 0.8]; (C) [0.1, 0.5] and [0.5, 0.9].

**SI Figure SI-2:**
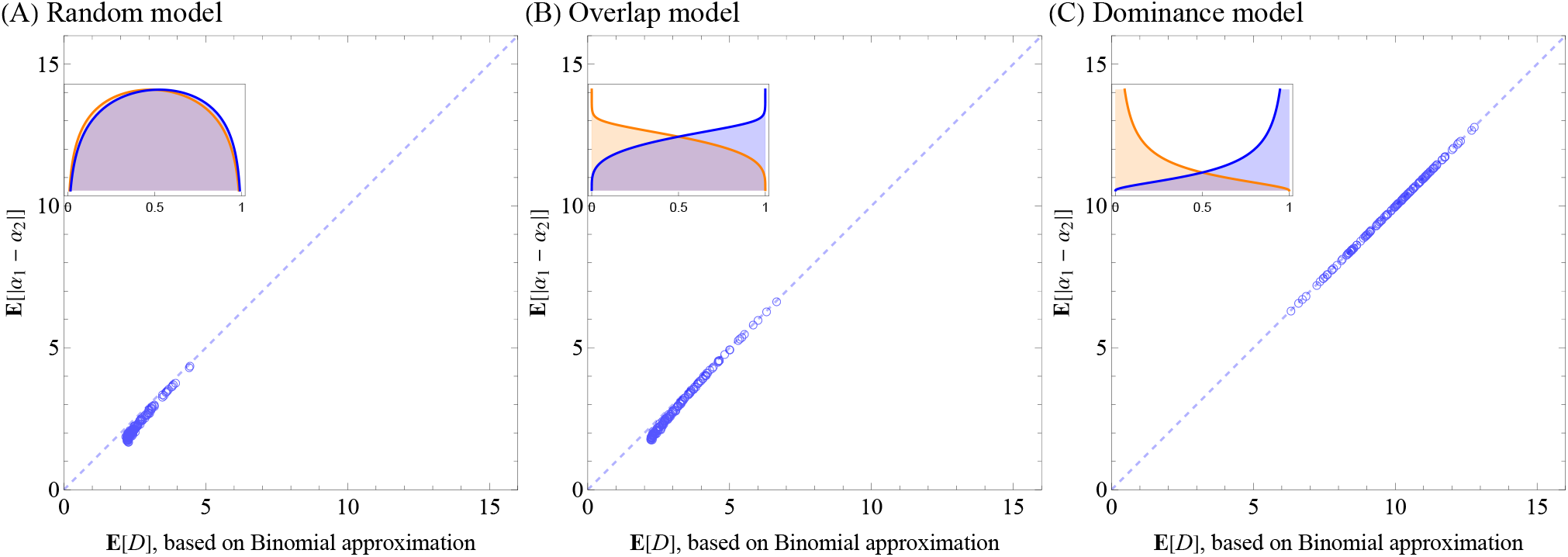
Approximation, based on the averaging method.

**SI Figure SI-3:**
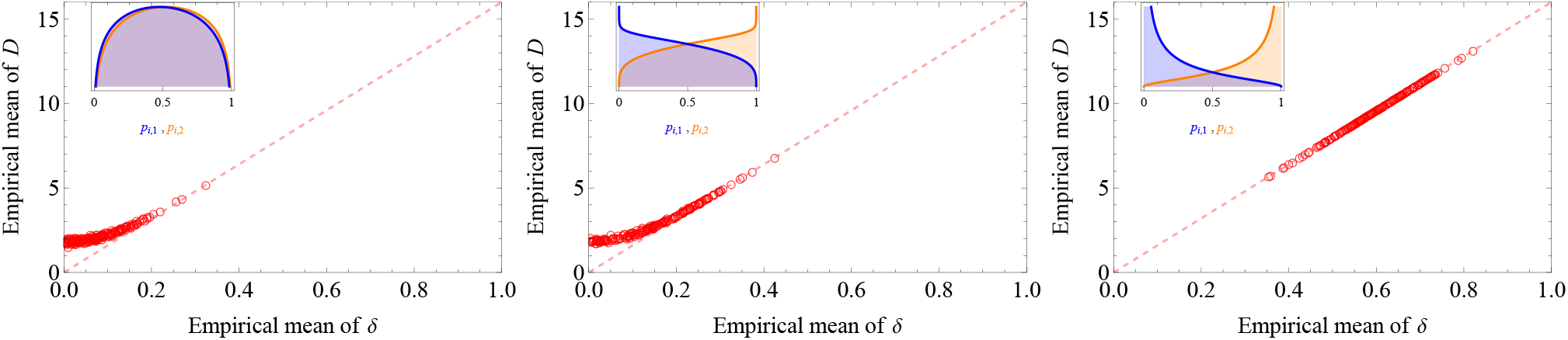
Plotting the relation between (i) abscissa: the average value of δ estimated from 300 incidence tables generated by a common probability distribution, versus (ii) ordinate: the average value of *D* from the 300 incidence tables generated by a common probability distribution. Each circle corresponds to a single simulation run (200 runs).

## Notes

### Competing Interest Statement

The authors have declared no competing interest.

### Summary of Updates

Some technical errors corrected.

